# Syndromic cholera diagnosis masks diverse causes of diarrhoeal disease in Burundi revealed by portable metagenomics

**DOI:** 10.64898/2026.03.23.713584

**Authors:** Emilie Egholm Bruun Jensen, Néhémie Nzoyikorera, Mirena Ivanova, Pimlapas Leekitcharoenphon, Marie Noelle Uwineza, Idrissa Diawara, Joseph Nyandwi, Frank M. Aarestrup, Saria Otani

**Affiliations:** Research group for Genomic Epidemiology, National Food Institute, Technical University of Denmark, 2800 Kgs Lyngby, Denmark; National Reference Laboratory, Institut National de Santé Publique (INSP) du Burundi, Bujumbura, Burundi; Research Laboratory of Microbiology, Infectious Diseases, Allergology and Pathogen Surveillance (LARMIAS), Mohammed VI Faculty of Medicine, Mohammed VI University of Sciences and Health (UM6SS), Casablanca, Morocco; Mohammed VI Higher Institute of Biosciences and Biotechnologies, Mohammed VI University of Sciences and Health (UM6SS), Casablanca, Morocco; Mohammed VI Center for Research and Innovation, Rabat, Morocco; National Public Health Institute, Ministry of Public Health Bujumbura, Burundi; Faculté de Médecine, Université du Burundi, Bujumbura, Burundi

## Abstract

**Background:** Cholera outbreaks remain a major public-health challenge in sub-Saharan Africa, where diagnostic capacity is limited and clinical case definitions are non-specific and reply heavily on syndromic diagnosis. Rapid identification of *Vibrio cholerae* is critical, yet cholera-suspected diarrhoea can have multiple infectious causes not captured by targeted diagnostics.

**Methods:** We evaluated a mobile, culture-independent metagenomic sequencing workflow for on-site detection of gastrointestinal pathogens directly from faecal samples in Burundi. The offline workflow combined long-read ONT sequencing with rapid, laptop-based taxonomic and antimicrobial resistance (AMR) screening and was deployed across a health centre, a district hospital, and a refugee transit camp. The frontline and real-time results were verified using both conventional culturing and in-depth bioinformatic analyses.

**Results:** *V. cholerae* signals were only detected in a subset of suspected cholera cases, while many samples were dominated by alternative bacterial taxa, most frequently *Escherichia coli*. *V. cholerae* abundance correlated strongly with detection of the cholera toxin phage CTXφ, supporting differentiation between toxigenic signal and background exposure. AMR genes were detected across samples, providing early situational insight into resistance determinants among gastrointestinal bacteria.

**Conclusions:** Mobile, offline metagenomic sequencing enables rapid frontline characterization of gastrointestinal disease, especially cholera-suspected, in resource-limited settings and complements existing diagnostics by improving etiological resolution and outbreak response.

**Author Summary:** Cholera remains a major cause of severe diarrhoeal disease in many low-resource settings, where diagnosis often relies on symptoms and limited laboratory testing. However, patients suspected of cholera can be infected by a wide range of other pathogens that are not detected by standard diagnostics. In this study, we evaluated a portable, sequencing-based approach that allows direct identification of pathogens from stool samples at the point of care, without the need for laboratory infrastructure, internet access, or culture.

Using this approach in multiple settings in Burundi, including a health centre, hospital, and refugee camp, we found a subset of suspected cholera cases were associated with *Vibrio cholerae*. Other cases were also dominated by other bacteria, particularly *Escherichia coli*. We also detected antimicrobial resistance genes across samples, providing additional information relevant for treatment and surveillance.

Our findings demonstrate that mobile metagenomic sequencing can improve the identification of disease causes directly in outbreak settings and help distinguish true cholera cases from other gastrointestinal infections. This approach has the potential to strengthen outbreak response, improve patient management, and support more accurate disease surveillance in resource-limited environments.

## Introduction

Acute gastrointestinal infections, particularly cholera, remain a recurrent and rapidly escalating cause of watery diarrhoea in many parts of sub-Saharan Africa.^1–3^ Outbreaks often emerge in settings with fragile water and sanitation infrastructure and can be amplified by displacement and constrained healthcare resources.

Because severe dehydration can develop within hours with diarrhoeal diseases,^4–5^ outbreak control and patient management depend on timely recognition of true cholera cases and differentiating them from other causes of gastrointestinal illness with overlapping symptoms. At the same time, confirmation frequently relies on centralised laboratory capacity and sample transport from remote places, and under-testing can delay targeted public-health action.

Despite its clinical and public-health importance, rapid and accurate identification of *Vibrio cholerae* remains challenging in many endemic settings.^1–3^ Standard and conventional culture requires selective media, time for incubation and large equipped laboratories that are often unavailable in peripheral health centres. Rapid diagnostic tests, while easier to deploy, are typically limited to detecting a narrow range of *V. cholerae* serogroups and do not capture other gastrointestinal pathogens that can cause clinically similar symptoms.^6–8^ Symptom-based diagnoses cannot reliably distinguish cholera from other causes of acute diarrhoeal disease,^9^ leading to under- or over-estimation of cases during outbreaks. In addition, this will not identify healthy carriers. These limitations compromise case management, delay outbreak confirmation needed for effective public-health response.

To address these diagnostic constraints, there is growing interest in approaches that can provide rapid, culture-independent identification of gastrointestinal infection agents,^10–13^ including *V. cholerae*, directly from clinical samples, while remaining deployable in resource-limited and outbreak-prone settings. An optimal frontline diagnostic workflow would operate without reliance on centralized laboratory infrastructure, cold-chain stability, continuous grid power, or internet connectivity, while still enabling broad detection of bacterial pathogens and preferably the associated antimicrobial resistance determinants. Recent advances in portable sequencing platforms have created new opportunities to move genomic diagnostics closer to the point of care, allowing real-time generation and analysis of pathogen data directly at outbreak sites and from the clinical samples without culturing.^10–14^ A portable and cost-effective sequencing method has been introduced with the MinION from Oxford Nanopore

Technologies (ONT).^15^ Over the years, the technology has improved significantly with the most recent MinION flow cell (R10.4.1) and the 14 chemistry library preparation have an increased accuracy of >99%.^16^ Together with the real-time basecalling, the portable laptop-compatible sequencing allows for a mobile laboratory setup, well-equipped for detection of infectious diseases directly at the outbreak site. With a metagenomic approach, where no culturing is needed, long-read sequencing can be used for frontline, on-site pathogen and antimicrobial resistance (AMR) detection.^15,17^ This can assist the healthcare professionals in treatment decisions and support timely public health interventions.

In Burundi, cholerae (*Vibrio cholerae*) is considered an endemic disease,^18^ but with occasional outbreaks associated with seasonality (the wet season creates conditions that promote the growth and survival of *V. cholerae*), inadequate sanitation, limited access to clean drinking water, and natural disasters.^1^ Epidemic cholerae is caused by two toxigenic serogroups O1 and O139,^19^ but O139 has so far not been isolated in Burundi.^18^ Similar to many countries in sub-Saharan Africa, Burundi faces several socioeconomic challenges that affect access to essential services, including healthcare, especially in rural and peri-urban areas.^20^ A large proportion of the population lives in rural areas with limited infrastructure, and ongoing humanitarian pressure from internal displacement and refugee movements (app. 89.000 refugees and asylum seekers so far)^21^ has further stretched already limited health resources.^22^ Access to specialised laboratory diagnostics is largely centralized, with comprehensive testing services typically only available in Bujumbura, making timely confirmation of infections difficult for peripheral clinics. Travel times from rural and camp settings to reference laboratories can be long and unpredictable creating barriers to both routine and outbreak-responsive diagnostics.

While portable sequencing has increasingly been applied in outbreak investigations, bioinformatic analyses are typically performed off-site due to the computational demands of large bacterial reference databases. In this study, we demonstrate that reducing bacterial reference database complexity while retaining species-level coverage enables rapid, fully offline pathogen detection directly in the field. We evaluated a mobile, culture-independent metagenomic workflow for on-site detection of *V. cholerae* and other gastrointestinal pathogens directly from faecal samples in Burundi. The workflow was designed to operate without grid power or internet access, enabling frontline diagnostics in resource-limited and outbreak-prone settings.

## Materials and Methods

### Sample collection and study design

In September 2025, field visits were conducted at Rugombo Health Centre, which hosts the Cholera Treatment Center (Figure 1A) to assess direct detection of *V. cholerae* in patient faecal samples. The workflow was subsequently evaluated at two additional sites: Rutana District Hospital, and Cishemere refugee camp in Rutana Province (Figure 1A) to confirm the workflow efficacy and its feasibility across distinct clinical settings and pathogens. At Rugombo Health Centre, 31 suspected cholera cases were investigated. On-site diagnostics was performed following our workflow (Figure 1B), including faecal sample collection, DNA extraction, library preparation, sequencing, (Graphics Processing Unit) GPU-enabled basecalling and rapid identification of pathogen and AMR locally using a lap-top based solution. Faecal samples from five asymptomatic companions (from the same household as the patients) from the Rugombo Health Centre were also collected and analysed in parallel. Additionally, to explore potential environmental sources and exposure routes in the Rugombo area, environmental samples from the Mparambo neighbourhood in Rugombo were collected. This included samples from a communal latrine, from an open drain, and river water which served both for bathing and for water collection for household purposes.

**Figure 1:**
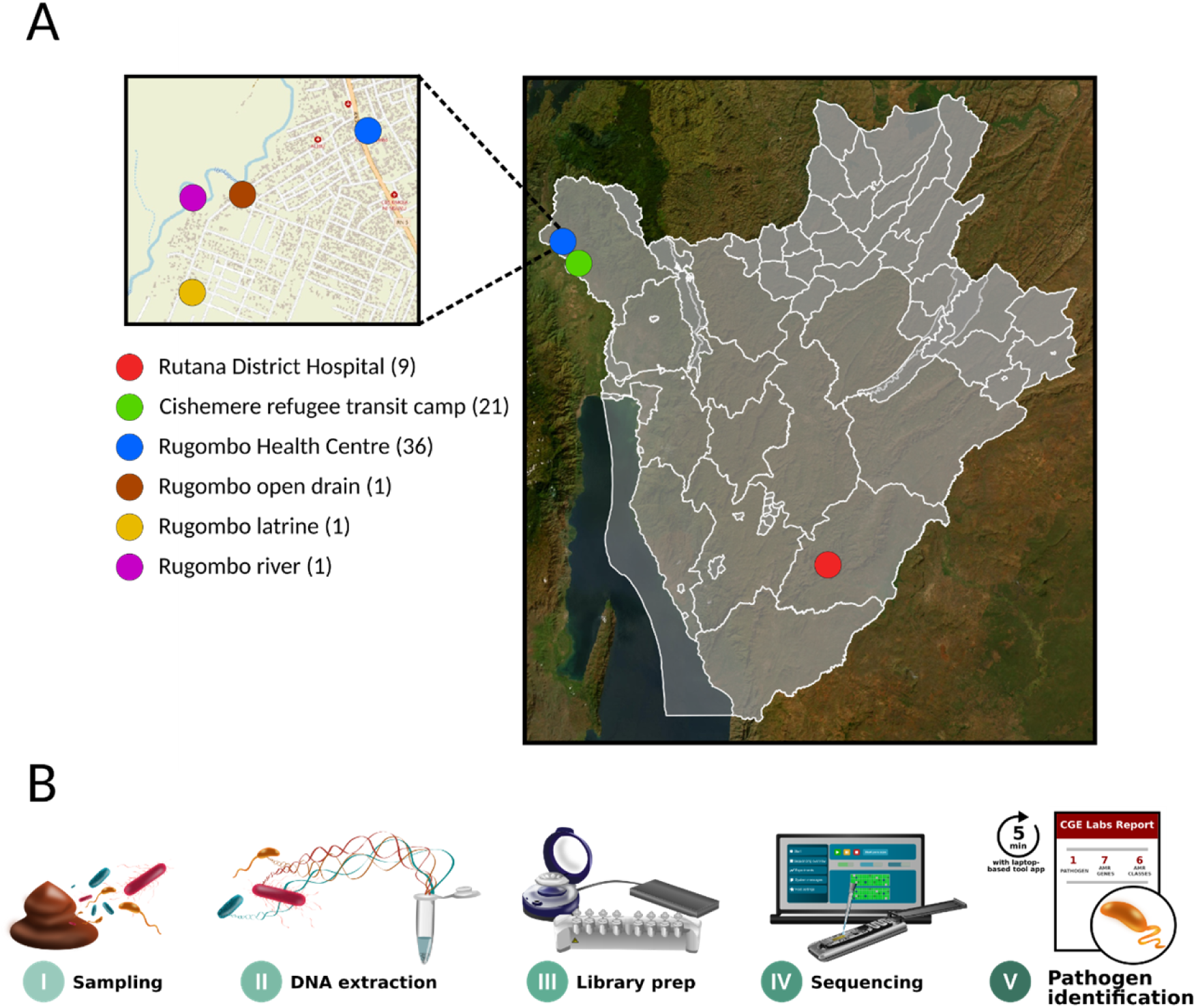
Study overview. **A)** Map showing the location of the metagenomic sample sites in Burundi. **B)** On-site rapid diagnostic workflow. This includes sample collection, extraction of DNA, library preparation, sequencing and basecalling, as well as pathogen and AMR identification using a local laptop-based solution. All conducted without access to grid power or internet.

The same setup was tested at the Cishemere refugee transit camp in Cibitoke (Figure 1). Here, 12 faecal samples were collected from individuals reporting stomach issues (discomfort with no clinical diagnosis), and nine from individuals without symptoms. At Rutana District Hospital, nine faecal samples were collected from admitted patients with gastrointestinal symptoms. Table S1 provides an overview of all samples analysed including the minimal metadata collected.

### ONT metagenomic sequencing

High molecular weight (HMW) DNA was extracted on-site from the faecal samples using Quick-DNA HMW Magbead Kit (Cat. No. D6060, Zymo Research) following the manufacturer’s instructions. The heatblock and centrifuge required for DNA extraction were powered using an external battery. The long-read library was prepared using the Ligation sequencing gDNA Native Barcoding Kit 24 V14 (SQK-NBD114.24, Oxford Nanopore Technologies). Enzymes and reagents which needed to be stored cold were kept on gel packs in a thermal container on-site. Samples were multiplexed and loaded onto the FLO-MIN114 flowcell (R10.4.1 chemistry). Sequencing was performed for a duration ranging from 20 to 45 hours, and the reads were basecalled on a GPU-enabled lap-top using the high-accuracy basecalling option (model dna_r10.4.1_e8.2_400bps_hac@v5.0.0) from Dorado version 0.9.6+49e25e9. Demultiplexing and default quality control (Q-score >9, and minimum read length >200 bp) was carried out in MinKNOW version 25.05.14. Sequencing output statistics were calculated with NanoStat version 1.6.0.^23^ and host reads were removed by Minimap2 version 2.28.^24^ alignment against the Hg38 reference genome (GCF_000001405.40).

### Local laptop-based pathogen and AMR detection

For rapid and on-site taxonomic and AMR screening, demultiplexed FASTQ reads were concatenated and mapped with KMA version 1.5.0.^25^ against a reduced-size bacterial reference database to enable the quick and offline analysis on the laptop. Database reduction was achieved by collapsing within-species strain diversity (i.e., selecting fewer representative genomes per species) while retaining species-level coverage. The reduced database contained 11,602 bacterial entries. AMR gene detection was performed using the entire AMR reference database ResFinder.^26^ The alignment with KMA was performed with a minimum identity threshold of 25% (-ID 25) and a minimum depth threshold of 1 (-md 1).

### Bacterial culturing and characterisation

To further confirm the presence of *V. cholerae* in the faecal samples from Rugombo, the samples were enriched in alkaline buffered peptone water (ABPW, pH 8.6 ± 0.2) prior to plating on thiosulphate citrate bile sucrose (TCBS) agar (Sigma) following the manufacturer’s instructions. After incubation, colonies displaying the expected colour (yellow) were selected as potential *V. cholerae* isolates. DNA was extracted and ONT libraries were prepared from those isolates as described before.^27^ Assembly was performed with Flye version 2.9.6 (--nano-hq) and Medaka version 2.0.1.^28^ was used to polish and create the consensus sequence (--model r1041_e82_400bps_hac_v5.0.0). One of the isolates (R24) had a sufficient high amount of reads for closing the genome. Trycycler version 0.5.4.^29^ was used to circularise the genome. Assemblies from two isolates (R14, R23) were binned with MetaBat2 version 2.17.^30^ to retrieve only pure and high completeness *V. cholerae* genomes. KmerFinder version 3.0.2 (database version: 2022-07-11)^25,31–32^ was used to verify taxonomy, and ST-type and serotypes were identified with MLST version 2.0.9 (database version: 2025-09-15),^33^ and CholeraeFinder version 1.0.^34^ The genomes were annotated with Bakta version 1.11.3 with database version 6.0.^35^ A phylogenetic tree based on core SNPs was created with CSI phylogeny version 1.4.^36^ with *V. cholerae* N16961 as reference genome (accession no. NZ_CP028827.1).

### Comprehensive HPC-based analyses

To complement the local on-site laptop-based detection of pathogens and AMR genes, a more elaborate command-line off-site approach was also applied using the HPC (high-performance computing) infrastructure at DTU. The concatenated FASTQ reads were mapped and aligned with KMA version 1.4.15 to three reference databases: The first database was a full (not reduced) custom reference genomic database created from NCBI GenBank (downloaded and created 05.24.2022) and similar to Osakunor et al.,^37^ and Otani and Jespersen et al.^38^ It consisted of completed and draft bacterial genomes. The sequencing reads were also mapped to a custom viral database created from NCBI GenBank (downloaded 04.12.2025). A template coverage threshold of >50% was used for the alignments. Additionally, a *V. cholerae*-specific database was created from the five Burundi isolates from this study, as well as 48 genomes from the *V. cholerae* isolate collection from PubMLST (downloaded 12.02.2026) which passed the following criteria: source = human, number of contigs < 10, total length (Mbp) > 3.8, and N50 > 50.000. The reference genomes were padded with 800 characters of N’s to optimise KMA conclave scoring for resolving multiple template matches and better distinguish between isolates.

### Data analysis and software

The kbio.math.diversity.alpha python package^39^ was used to describe alpha diversity by calculating richness (chao1), Simpson diversity index, and Shannon’ evenness. Beta-diversity was described with a compositional data analysis approach^40^. With python, the package pyCoDaMath^41^ was used to perform zero replacement and transform the abundance tables to CLR-values. The same package was used to create the PCA figures.

Differential abundance analysis was performed with ALDEx2 version 1.42.0.^42^ to identify differential bacterial species between groups. With a Welch’s *t*-test followed by a Benjamini-Hochberg false-discovery rate (FDR) correction, differences in the CLR abundances between groups are identified. Bacterial species with absolute effect size >0.8 and FDR <0.05 were reported as statistically significant for each group. The effect size was calculated as seen in equation 1.

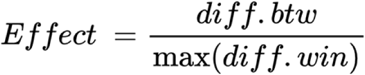

**Equation 1**: Calculation of effect size used in the differential abundant analysis. Effect is the median difference in CLR values between groups (diff.btw) divided by the median of the largest difference in CLR values within groups (diff.win).

Linear association between the abundance of bacteria and viruses was tested using Pearson’s correlation coefficient with the python SciPy library.^43^ The basepair counts were normalised to adjust for reference length and bacterial depth using equation 2.

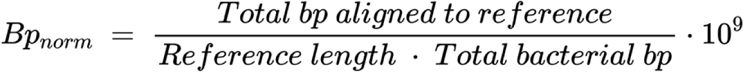

**Equation 2**: Basepair normalisation to account for reference length and bacterial depth.

## Results

### Sequencing output

Five FLO-MIN114 flowcells (R10.4.1 chemistry) were used for the metagenomic sequencing (Figure 2). The samples from Rugombo Health Centre were sequenced on two flowcells (based on the flowcell availability at that site at the moment), with 12 and 24 samples for 45 hours and 7 minutes, and 22 hours and 40 minutes, respectively. The flowcell with 12 samples yielded 3.97 giga bases (Gb), 1.2 M reads, a median quality of Q16.35, and a N50 of 4.93 kilo basepairs (bp). The second flowcell from Rugombo Health Centre with 24 samples yielded 6.83 Gb, 3.2 M reads, a median quality of Q15.10, and a N50 of 3.36 kb. The 21 samples from Cishemere refugee camp were sequenced on two flowcells for 32 hours and 33 minutes, and 10 hours and 55 minutes respectively. The first flowcell yielded 0.52 Gb, 0.2 M reads, a median quality of Q15.55, and a N50 of 4.62 kb. The second flowcell yielded 5.55 Gb, 2.2 M reads, a median quality of Q15.5, and a N50 of 5.07 kb. The nine samples from Rutana District Hospital and the three environmental samples from the Mparambo neighbourhood in Rugombo were multiplexed and sequenced on one flowcell for 24 hours and 53 minutes, and yielding 2.28 Gb, 0.9 M reads, a median quality of Q13.65, and a N50 of 5.26 kb. All flowcell outputs are summarised in Table S2 as well.

**Figure 2:**
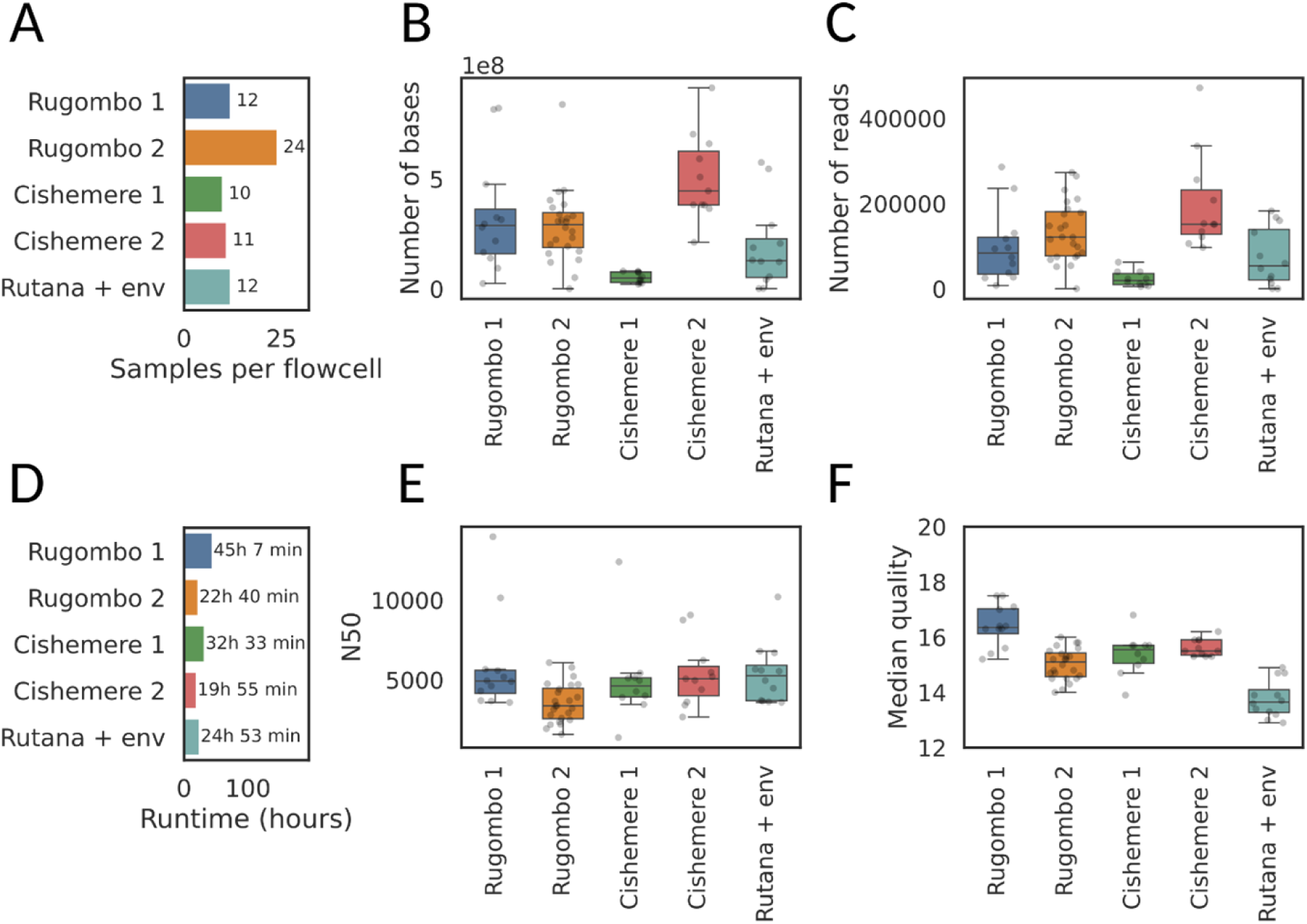
Sequencing output summary of the five flowcells with faecal samples. **A)** Number of samples barcoded per flowcell. **B)** Number of bases sequenced per sample per flowcell. **C)** Number of reads per sample per flowcell. **D)** Sequencing time. **E)** N50 per sample per flowcell **F)** Median quality per sample per flowcell.

### Metagenomic mobile laboratory on-site diagnostics of *V. cholerae* at Rugombo Health Centre

*V. cholerae* was detected in high abundance in 11 of the 31 patient samples (Figure 3). In six of these *V. cholerae*-positive cases, high abundance of *E. coli* was also observed, and in one of the samples (R09) *V. cholerae*, *E. coli*, and *Campylobacter jejuni* were detected (Figure 3A), with *C. jejuni* being low abundant (i.e. 0.976% of the sequenced bacterial basepairs) Nine samples had *E. coli* as the only potential pathogen detected (Figure 3A). These lap-top based results were subsequently confirmed by alignment to the full comprehensive genomic database (Figure 3A). Two samples (OR_20 and R22) also had high abundance of *Candidatus Campylobacter infans* (Figure 3A), which has been shown to be present in both symptomatic and asymptomatic children under two years with diarrhoea,^44^ and is therefore not regarded as a causative agent. No apparent potential pathogens were detected in the remaining 10 patient samples (Figure 3A) However, remaining non-pathogenic African microbiome members^38^ were detected in all the samples, e.g., *Bacteroides*, *Blautia*, *Roseburia* and *Prevotella* (Figure 3). No pathogens were detected in any of the asymptomatic companions. The gut microbiome of these individuals were characterised by a high abundance of *Klebsiella quasipneumoniae*, formerly named *Prevotella*-species like *Segatella hominis* (formerly *Prevotella hominis*) and *Leyella stercorea* (formerly *Prevotella stercorea*), and *Escherichia coli* (Table S3). Faecal indicator bacteria such as *Klebsiella quasipneumoniae*, *S. hominis*, and *E. coli* were also found in high abundance in the open drain and river environmental samples (Table S4, Figure 3A). The latrine microbiome showed high similarity to the asymptomatic companions, whereas the river and open drain sample clustered within the patient samples (Figure S1A).

**Figure 3:**
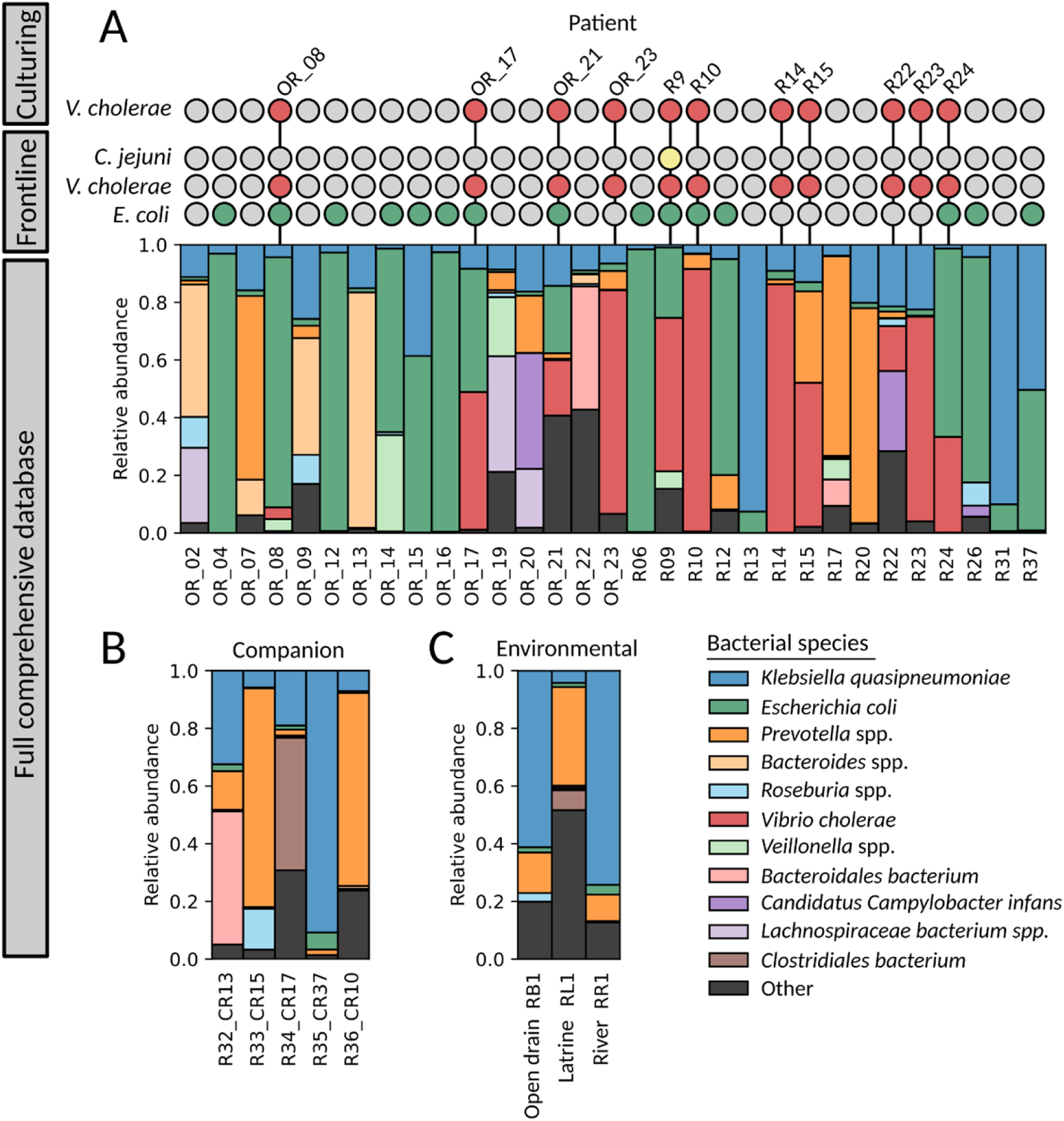
Metagenomic insight into the faecal samples collected from patients with *Vibrio cholerae*-suspected infections as well as companions- and environmental samples from Rugombo Health Centre. **A)** Comparison between culturing on TCBS plate agar for *V. cholerae* detection, frontline laptop-based diagnostics results with a reduced database, and the command line with a comprehensive database. The result shows a presence/absence of the pathogen from the culturing and frontline laptop-based alignment. Red indicates detection of *V. cholerae*, green indicates *E. coli*, and yellow indicates *C. jejuni*. Underneath, the barplot shows the in-depth compositional analysis of the gut microbiome obtained from the alignment to the full comprehensive database. **B)** Gut microbiome composition of the asymptomatic companions from the alignment to the full comprehensive database. No pathogens were detected. **C)** Bacterial composition of the environmental samples from the alignment to the full comprehensive database. No pathogens were detected.

The gut microbiome of the companions had a higher richness than the patients (Figure S1B). Differential abundance analysis comparing the 10 cholerae-negative patients with the asymptomatic companions found four bacterial species significantly more abundant in the asymptomatic companions, while none was found in the cholerae-negative patients (Figure S1C): *S. hominis*, *Thalassococcus profundi*, *Alcanivorax profundi*, and *Prevotella pectinovra*. Only *V. cholerae* were found significantly more abundant in the cholerae-positive samples when compared to the asymptomatic companions (Figure S1D).

AMR genes identified in patients included resistance to several antimicrobial classes, with the following being the most dominant: beta-lactams, tetracyclines, macrolides, and quinolone (Figure S2B). Additional alignment to a reference virus database was performed. However, none of the patient samples had reads mapping to common diarrheal-causing viruses such as adenoviruses (Figure S3).

### Isolation of *V. cholerae* from Rugombo Health Centre

Presence of *V. cholerae* was supported by bacterial culturing from the faecal sample. The 11 faecal samples positive of *V. cholerae* from the on-site alignment also showed morphologically positive *V. cholerae* colonies on TCBS agar (yellow colonies). Those were OR_08, OR17, OR_21, OR_23, R9, R10, R14, R15, R22, R23 and R24. All of those isolates except one (OR_17) were also whole genome sequenced (WGS) with long reads. Five of which were identified as pure *V. cholerae* (Table 1), while the remaining five isolates were identified as a mixture of different taxa, yet *V. cholerae* was the most abundant in the culture. Four of the isolates were ST69 with O1 and El Tor variant, and clustered together with a difference of 0-20 SNPs (Figure S4). The non O1/O139 strain (R24) was circularised, and as expected, it clustered separately from the four other isolates (Figure S4).

**Table 1:** Characteristics of the five *V. cholerae* strains.

| Sample | No. of contigs | Total length (bp) | GC content (%) | ST type | Serogroup |
| --- | --- | --- | --- | --- | --- |
| R15 | 6 | 4,232,353 | 0.476 | ST69 | O1/El Tor variant |
| R10 | 3 | 4,224,272 | 0.476 | ST69 | O1/El Tor variant |
| R14 | 2 | 4,085,442 | 0.475 | ST69 | O1/El Tor variant |
| R23 | 60 | 3,937,251 | 0.476 | ST69 | O1/El Tor variant |
| R24 | 2 | 4,170,543 | 0.476 | ST366 | Non-O1/non-O139 |

To identify which *V. cholerae* strains were present in the metagenomic samples, a database of *V. cholerae* genomes was created from downloaded reference genomes of *V. cholerae*, as well as the five isolates from this study (see methods). Sequence alignment to this cholerae-specific database confirmed *V. cholerae* presence in the 11 cholerae-positive samples previously identified (Table S5). For all 11 samples, O1/El Tor variant from Burundi 2025 was the dominant one, while R24 had a mix of the O1/El Tor variant and the non-O1/non-O139, with the latter being in agreement with our WGS result.

### Pipeline validation at additional sites

Two additional sites were also visited: Rutana District Hospital and Cishemere refugee transit camp in Cibitoke. From Rutana District Hospital, nine faecal samples were collected from patients with gastrointestinal symptoms. *E. coli* was identified as the pathogen in five of the nine samples, and this was confirmed with the command-line alignment to the comprehensive genomic database (Figure 3C, Table S6). In three of the four samples where *E. coli* was not the dominant species (RH3, RH6, and RH9), *Segatella copri* was found abundant. AMR genes identified in the Rutana samples included resistance to e.g. Beta-lactams (all samples), tetracycline (all samples), macrolides, and quinolones (Figure S2A).

On-site at the Cishemere refugee camp in Cibitoke, 12 individuals who reported stomach pain or discomfort were investigated. The faeces appeared compact and showed no signs indicating diarrhoea. Additional nine samples were also collected from nine asymptomatic individuals. No pathogens were detected using our on-site sequencing pipeline for any of the samples (Figure 4A). The in-depth analysis of the bacterial composition found no reads mapping to either *V. cholerae* or *C. jejuni*, and no difference between the individuals with stomach discomfort and asymptomatic individuals were identified either (Figure 4B).

**Figure 4:**
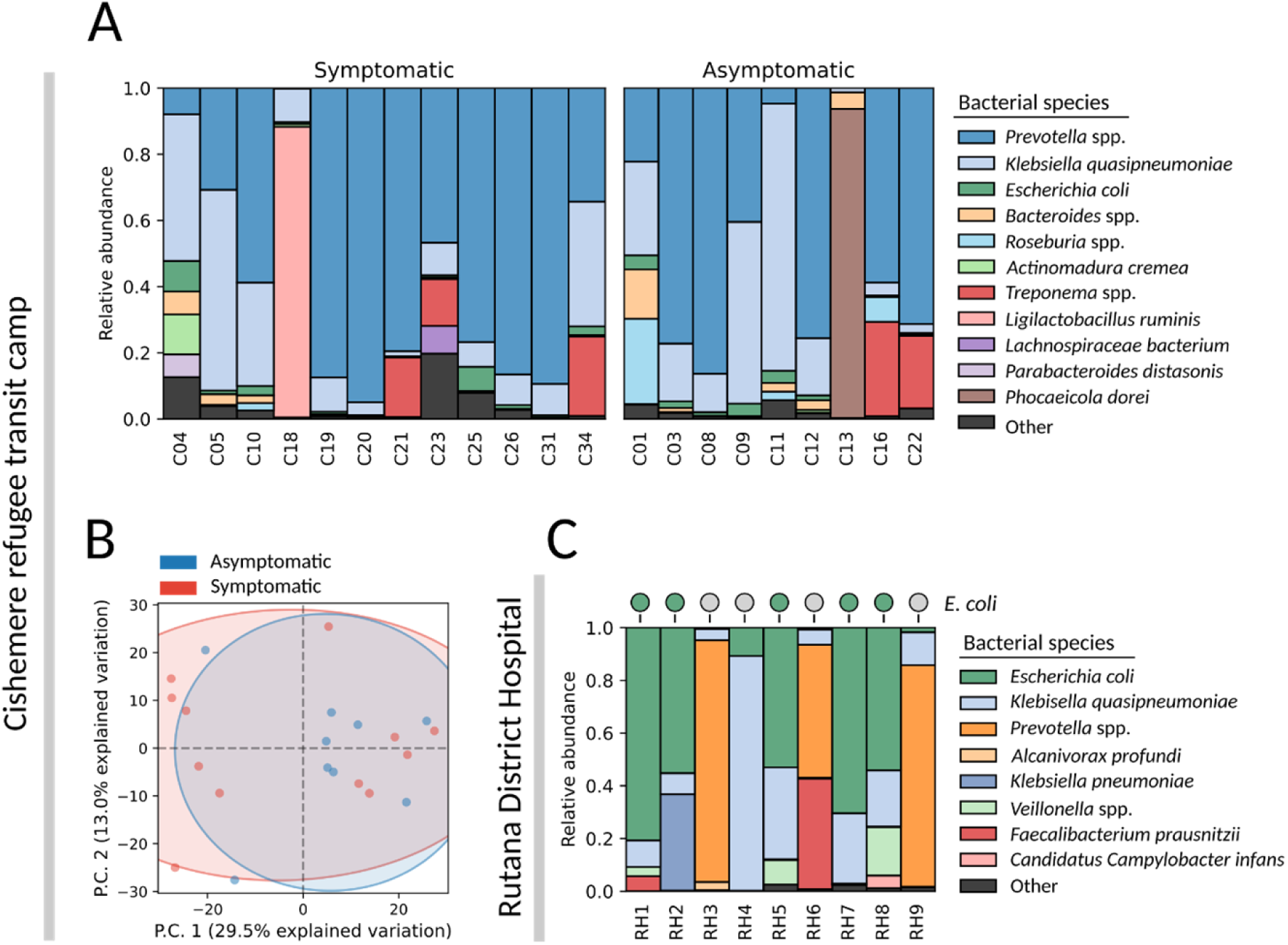
Metagenomic insight into the faecal samples collected from individuals with stomach discomfort and from asymptomatic individuals at the Cishemere refugee camp in Cibitoke, as well as from patients with suspected gastrointestinal infections at Rutana District Hospital. **A)** Bacterial species composition of the faecal samples collected from individuals at the Cishemere refugee transit camp. **B)** PCA clustering of the samples from Cishemere refugee camp based on the bacterial species. **C)** Comparison of the frontline diagnostic results at Rutana District Hospital obtained from the lap-top based alignment against a reduced bacterial database with the in-depth command-line based approach with a comprehensive bacterial database. The result shows a presence/absence of the pathogen from the on-site diagnostics, as well as a barplot showing the abundance of bacterial species within the sample from the command-line approach.

The most abundant bacterial taxa in both symptomatic and asymptomatic individuals were *S. hominis*, *K. quasipneumoniae*, and *Leyella stercorea* (previously *Prevotella stercorea*) (Table S7). Other dominant species were *Treponema* spp. and *Roseburia* spp. (Figure 4A), both reported common members of the sub-Saharan human gut microbiome.^38^ Antimicrobial resistance patterns were identified within the samples: All individuals with stomach discomfort harboured genes encoding resistance to nitroimidazoles, beta-lactams, tetracyclines, and nitroimidazoles (Figure S2C). This resistance profile was also observed for most of the asymptomatic individuals. Additional alignment to a virus reference database (see methods) was performed to determine if the symptomatic individuals had a viral infection such as adenoviruses. However, no reads aligned to DNA viruses. (Figure S3C).

### Presence of CTXφ confirms pathogenicity of *V. cholerae*

Investigations into the abundance correlation between bacteriophages and their bacterial hosts spanning all samples across all sites, revealed that the abundance of *V. cholerae* was positively correlated with the abundance of the filamentous bacteriophage Cholera Toxin Phage (CTXφ) (Figure S5) with a Pearson correlation coefficient of 0.851. The pathogenicity of *V. cholera* often comes from the integration of the Cholera Toxin Phage (CTXφ) into the genome.^45^ The *E. coli* phage phiSTEC1575-Stx2k as well as the *E. coli* phage RCS47 were both found linearly associated with the abundance of *E. coli*, with a Pearson correlation coefficient of 0.850 and 0.498, respectively. The Stx2k phage carries Shiga toxin type 2 and is able to infect and integrate the gene into the genome, which convert non-pathogenic *E. coli* into a highly pathogenic and Shiga toxin-producing strain (STEC)^46^. The *E. coli*-infecting RCS47 P1-like bacteriophage have been reported in clinical *E. coli* strains and are associated with the beta-lactam resistance gene *bla*SHV-2.^47^

## Discussion

Here, we evaluated a fully mobile, offline ONT metagenomic workflow for rapid on-site detection of *V. cholerae* and other gastrointestinal pathogens directly from faecal samples in Burundi. The workflow was designed for outbreak-prone, resource-limited settings without grid power or internet access, and combines rapid, laptop-based screening with subsequent in-depth read mapping for confirmation. Across three distinct settings: A health centre evaluating suspected cholera cases, a district hospital, and a refugee transit camp, we demonstrate that culture-independent long-read metagenomics can generate actionable pathogen profiles on-site, in approximately 12 hours from sample to report, and reveal substantial heterogeneity in the aetiology of “cholera-suspected” gastrointestinal disease.

Conventional diagnostic methods, including rapid antigen tests targeting *V. cholerae* O1/O139, remain valuable tools for frontline cholera detection. However, when rapid tests are negative, inconclusive, or when patients do not respond to treatment, untargeted metagenomic sequencing provides an important complementary approach by enabling broader detection of gastrointestinal pathogens without prior assumptions. In this context, ONT-based metagenomics can function as an escalation tool rather than a replacement for existing diagnostics, especially here where at least one of the *V. cholerae* cases was not O1/O139.

Only a subset of clinically suspected cholera cases showed high-abundance *V. cholerae* signals consistent with a causative role in such cases of diarrhoea,^48–49^ while a large fraction of samples were dominated by other taxa, most frequently *E. coli*. This is important for outbreak settings where symptom-based diagnosis is unavoidable but non-specific: Acute watery diarrhoea is clinically overlapping across multiple pathogens, and reliance on syndromic case definitions alone can both inflate cholera case counts and miss alternative causes requiring different clinical and public-health attention.^6–8^ By providing an empirical profile from a single assay, metagenomics can support a “syndromic” diagnostic approach that is particularly relevant when conventional testing is unavailable or focused on a single target.

Detection does not necessarily imply causation, a challenge for metagenomic frontline diagnostics. In this study, applying a template coverage threshold of >50% removed low-abundance *V. cholerae* mappings and restricted detection to samples where the bacterium dominated the microbial signal and aligned with the clinical picture of suspected cholera. This highlights the importance of implementing explicit interpretation criteria when using metagenomics for pathogen detection directly from clinical samples. In clinical metagenomics, low-level sequence detection can arise from colonisation, transient exposure, or environmental contamination rather than representing the primary aetiology of disease.^50^ For frontline detection pipelines, these findings support the use of abundance and coverage thresholds, contextual co-detections, and clinical plausibility rather than binary positive/negative reporting.

Metagenomic profiling further revealed that cholera-positive patients exhibited reduced bacterial diversity compared to asymptomatic companions, consistent with previous observations that *V. cholerae* infection is associated with a collapsed gut microbiome.^49,51^ *V. cholerae* was the only bacterial species enriched in cholera-positive samples (Figure S1D), indicating that the infection signal was largely driven by a single dominant pathogen rather than multiple co-occurring taxa. In contrast, four bacterial species, *Segatella hominis*, *Thalassococcus profundi*, *Alcanivorax profundi*, and *Prevotella pectinovora*, were more abundant in asymptomatic companions compared with cholera-negative patients (Figure S1C), suggesting that these taxa may characterize the background gut microbiome in non-cholera individuals.

Using long-read metagenomics on-site captures not only taxonomic signals, but also pathogenicity-derived markers that can further support our interpretation. In our dataset, the abundance of *V. cholerae* correlated strongly with CTXφ abundance, providing internal biological support that high-abundance detections reflected toxigenic cholera-associated signals rather than low-level background. The *ctx* genes are carried by CTXφ and are central to epidemic cholera,^45^ and incorporation of virulence gene evidence alongside species detection is a logical next step for metagenomic cholera confirmation directly from stool. More broadly, this illustrates how metagenomics can move beyond identifying taxa to supporting interpretation using biological context.

We also performed culture-based confirmation on a subset of samples and observed that selective enrichment and plating yielded presumptive colonies, but isolate purity remained variable, with WGS revealing mixed signals even from isolated colonies. This likely reflects the complexity of faecal material and the selectivity of differential media in field settings. This also supports a pragmatic view of culture confirmation during outbreak response: While culturing remains essential for phenotypic testing and downstream characterization, it can be slow and operationally constrained on-site, and it does not always provide immediate clarity at the point of care. Culture-independent sequencing can therefore complement culture by accelerating confirmation and guiding which samples should be prioritized for isolation, storage, and phenotypic work.

Several commensal taxa were more abundant in asymptomatic companions, including *Segatella hominis* and *Prevotella pectinova*. Previous studies have shown other commensal bacteria such as *Blautia* species can contribute to colonisation resistance against *V. cholerae* through interference with quorum sensing and degradation of host-derived virulence-inducing compounds.^49,52^ However, no such evidence exists in the case of *S. hominis* or *P. pectinova*. Short-chain fatty acid-producing bacteria do however provide colonization resistance against *V. cholerae*,^53–55^ and it can be speculated that the SCFAs produced by *S. hominis*^56–57^ and *P. pectinovora*^58^ provide similar protection. However further investigations are needed to confirm this.

Environmental sampling did not yield clear *V. cholerae* detections in our limited set of samples. However, the environmental metagenomes contained high abundances of fecal-associated taxa and potential indicator species, consistent with exposure to human waste. Environmental *V. cholerae* detection is inherently challenging because shedding is intermittent, pathogen abundance may be low relative to complex background communities, and sampling is sensitive to timing and hydrological conditions.^59^ The absence of detections could be difficult, particularly given the small number of samples and field constraints on access. Future work should combine repeated sampling with concentration strategies (e.g., filtration and enrichment) to increase sensitivity and improve the utility of metagenomic environmental testing for source attribution.

Our offline analysis implemented rapid on-site mapping using KMA against a reduced-size bacterial reference database to enable fast processing on a laptop, followed by more comprehensive mapping for validation. This multilayered approach aligns with the reality of outbreak response, where fast initial answers are needed for awareness, while deeper analyses can follow once time and compute allow. Such designs are increasingly recognized as critical to the successful integration of portable genomics into routine surveillance and response workflows.

Antimicrobial resistance signals were also detected (offline) across sites, including genes associated with multiple antibiotic classes. Genotype-based resistance prediction from stool metagenomes can provide early warning of resistance genes circulating in the local gut microbial reservoir. In cholera management, rehydration remains primary,^19^ but antibiotics may be used for severe cases and to reduce duration and shedding; therefore, resistance surveillance remains relevant to both clinical guidance and public-health planning. Our findings also show that a substantial fraction of cholera-suspected cases were dominated by other bacterial pathogens, e.g., *E. coli*, for which antimicrobial treatment is more commonly considered. In this, rapid AMR screening adds additional clinical and public-health value by providing early situational awareness of resistance determinants circulating among gastrointestinal pathogens beyond *V. cholerae*.

Future work should focus on our identified limitations during our on-site investigation. First, clinical metadata were minimal and we did not perform a prospective detailed comparison against serotype testing and culture for all samples; future evaluations should quantify sensitivity/specificity and formalize interpretation thresholds for *V. cholerae* and key GI pathogens in field conditions. Second, environmental sampling was limited and should be expanded with repeated sampling and concentration to enable stronger inferences about potential sources.

Our results support mobile, offline ONT metagenomic sequencing as a feasible frontline diagnostic approach for cholera-suspected gastrointestinal disease in resource-limited settings.

By enabling rapid, culture-independent detection of *V. cholerae* alongside alternative aetiologies and AMR signals, this workflow can strengthen outbreak confirmation, improve diagnostic specificity at the point of care and front-line responses, and support timely public-health intervention where conventional diagnostics are delayed or unavailable.

## Acknowledgements

We are thankful to Frederik Duus Møller, Nikiforos Pyrounakis, and Philip Thomas Lanken Conradsen Clausen for technical assistance.

## Data availability statement

The sequencing data generated during the study is available through the European Nucleotide Archive (ENA) repository: PRJEB108124. This includes metagenomic sequencing reads as well as reads and assemblies of five *V. cholerae* single isolates, including the circularised non O1/O139 strain (R24).

## Ethics approval

The ethical approval to conduct this study was given by the National Ethics Committee in Burundi (Reference number: CNE/10/2024). The national ethics committee provided a waiver for the consent from participants in this study. The collection of the faecal samples is non-invasive and were collected by health professionals at all sites.

## Author contribution

EEBJ, NN, MNU, ID, SO: Experimental setup, methodology, sampling strategy, sample collection from the field, DNA extraction and bacterial culturing. EEBJ, MI, PL and SO: Library preparation and sequencing and sequence analyses. EEBJ and NN: Data curation, validation and formal analysis. EEBJ, NN, FMA and SO: wrote the first draft. EEBJ, NN, MI, PL, JN, FMA and SO: Review & editing. NN, JN, FMA and SO: Project administration, resources, and funding acquisition.

